# Systematic analysis of mutational spectra associated with DNA repair deficiency in *C. elegans*

**DOI:** 10.1101/2020.06.04.133306

**Authors:** B Meier, NV Volkova, Y Hong, S Bertolini, V González-Huici, T Petrova, S Boulton, PJ Campbell, M Gerstung, A Gartner

## Abstract

Genome integrity is particularly important in germ cells to faithfully preserve genetic information across generations. As yet little is known about the contribution of various DNA repair pathways to prevent mutagenesis. Using the *C. elegans* model we analyse mutational spectra that arise in wild-type and 61 DNA repair and DNA damage response mutants cultivated over multiple generations. Overall, 44% of lines show >2-fold increased mutagenesis with a broad spectrum of mutational outcomes including changes in single or multiple types of base substitutions induced by defects in base excision or nucleotide excision repair, or elevated levels of 50-400 bp deletions in translesion polymerase mutants *rev-3*(pol ζ) and *polh-1*(pol η). Mutational signatures associated with defective homologous recombination fall into two classes: 1) mutants lacking *brc-1/BRCA1* or *rad-51*/RAD51 paralogs show elevated base substitutions, indels and structural variants, while 2) deficiency for MUS-81/MUS81 and SLX-1/SLX1 nucleases, and HIM-6/BLM, HELQ-1/HELQ and RTEL-1/RTEL1 helicases primarily cause structural variants. Genome-wide investigation of mutagenesis patterns identified elevated rates of tandem duplications often associated with inverted repeats in *helq-1* mutants, and a unique pattern of ‘translocation’ events involving homeologous sequences in *rip-1* paralog mutants. *atm-1/*ATM DNA damage checkpoint mutants harboured complex structural variants enriched in subtelomeric regions, and chromosome end-to-end fusions. Finally, while inactivation of the *p53*-like gene *cep-1* did not affect mutagenesis, combined *brc-1 cep-1* deficiency displayed increased, locally clustered mutagenesis. In summary, we provide a global view of how DNA repair pathways prevent germ cell mutagenesis.

## Introduction

Germ cells are required to pass genetic information from one generation to the next, rendering the maintenance of their genetic integrity particularly important. While germ cell mutations are the basis of evolution, mutational events tend to be detrimental and are associated with reduced fitness and inherited disease.

Endogenous mutagenesis can be caused by nucleotide mis-incorporation during replication and by reactive cellular metabolites. Hydrolytic reactions trigger abundant depurination and depyrimidination events, and the deamination of cytosine and 5-methylcytosine (for review (Lindahl and Barnes 2000)). Reactive oxygen species, byproducts of oxidative phosphorylation and oxygen-dependent enzymatic processes, induce 10,000-100,000 DNA lesions per cell per day including base modifications such as 8-oxo-dG, thymine glycol and DNA single-strand breaks (Ames et al. 1993). In addition, enzymatic and non-enzymatic mechanisms lead to base methylations. For instance, 3-methyl-adenine and 3-methyl-cytosine can lead to mutations by blocking replication and *O*^*6*^-methyl-guanine leads to G>A changes (for review (Lindahl and Barnes 2000)). Metabolic byproducts such as reactive aldehydes form DNA adducts that can crosslink bases on two complementary DNA strands generating obstacles to replication and transcription.

DNA double-strand breaks (DSBs) are one of the most toxic DNA lesions and arise when the replication fork is stalled by base modifications, repetitive DNA, DNA sequences prone to form tertiary structures, or collision with the transcription machinery (Mehta and Haber 2014). Nevertheless, some cellular events require DSBs to be induced naturally, for example during immunoglobulin gene rearrangement to ensure immunoglobulin diversification. Additionally, during germ cell meiosis multiple DSBs are introduced in each chromosome resulting in at least one crossover recombination event per chromosome to facilitate the exchange of genetic information and ordered chromosome segregation (for review (Hillers et al. 2017)). Recombination requires a free DNA end to search for and invade a homologous DNA strand which acts as a template to facilitate the restoration of genetic information. When DSBs occur in repetitive DNA such as tandem repeats or interspersed repeat elements like Line and Alu sequences, ‘homology search’ provides a formidable challenge.

Nevertheless, only a vanishingly small fraction of primary lesions leads to the formation of mutations, highlighting how effective DNA damage response mechanisms are in detecting and mending multifarious forms of DNA damage. The analysis of the observed mutations has the potential to shed light on the primary mutagenic lesion. When the amount of DNA damage introduced by a mutagenic process exceeds the capacity of DNA repair, distinct patterns of mutations, referred to as mutational signatures or spectra, arise. Here we characterize genome-wide mutational spectra by analysing the number and distribution of single and multi-nucleotide variants (SNVs and MNVs), small insertions and deletions (indels) and structural variants (SVs) composed of larger deletions, inversions, duplications and chromosomal translocations.

The rates and patterns of germline mutations were previously studied in humans and model organisms such as mouse, fruit fly, *C. elegans* and primates (Keightley et al. 2014; Rahbari et al. 2016; Venn et al. 2014; Denver et al. 2009). However, large-scale investigation of germline mutagenesis is highly time- and labour-intensive. *C. elegans* offers a suitable system to study mutation accumulation across multiple generations and genetic backgrounds based on its short life-cycle and its ability to self-fertilize. In our accumulation experiments (MA), we propagated clonal lines over up to 40 generations (**Fig. 1a**). Crucially, these lines pass through a single-cell bottleneck provided by the zygote, as part of each of the 3-4 days life cycle of the nematode, enabling us to analyse how mutations arise in the germline (**Fig. 1a**).

**Fig. 1.**
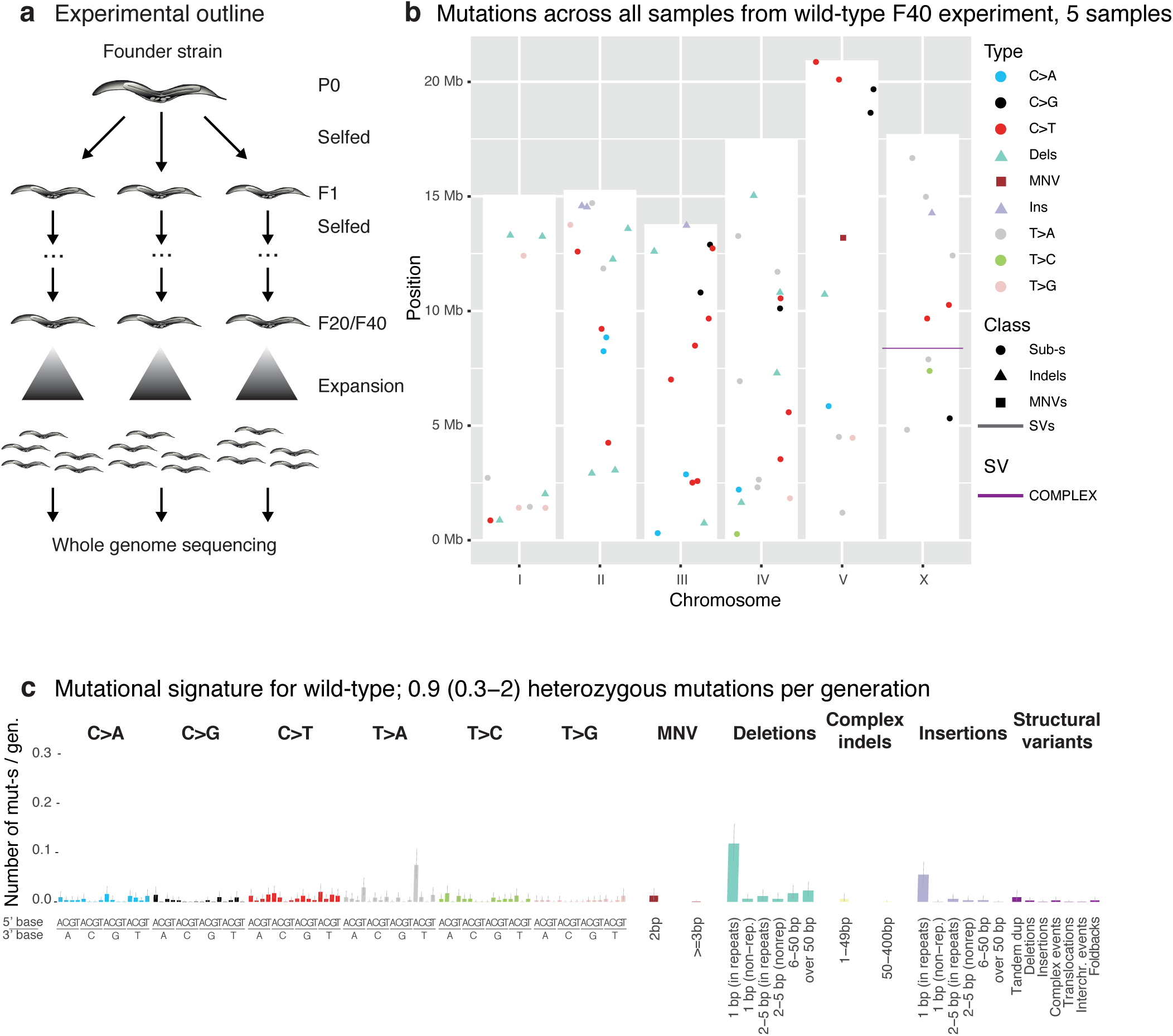
Experimental outline and background mutagenesis in wild-type. **a**. L4 larvae (F1 generation) from a parental founder strain (P0) were individually picked onto NGM plates and allowed to self-fertilize prior to picking individual L4 larvae of the next generation (F2) from each F1 plate. This process was repeated till clonal lines reach generation F20 or F40. Clonal lines were then allowed to expand, harvested, and prepared for whole genome sequencing (Methods). **b**. Mutation types and their location on the 6 *C. elegans* chromosomes (I-V and X) across all wild-type samples encompassing all mutation classes. The height of the white bars corresponds to the length of the respective *C. elegans* chromosome. Single nucleotide variants are indicated by a dot, dinucleotide variants (DNVs) by a square, indels divided in deletions (D) and insertions (I) by a triangle, and structural variants (SVs) by a line. **c**. Number of heterozygous mutations per generation across all mutation classes and types in N2 wild-type. Single nucleotide variants are shown in their 5’ and 3’ base context.

In this study, which encompasses wild-type, 54 single, and 7 double and triple mutants affecting all conserved DNA repair and DNA damage response pathways (**Table S1)**, we systematically determine the consequences of DNA repair and damage response deficiencies in the germ cell lineage and investigate the roles of various pathways in counteracting endogenous DNA damage.

## Results

### Mutation rates in wild-type and DNA repair deficient *C. elegans*

We employed 30 mutation accumulation (MA) lines, five of which were grown for 40 generations (**Fig. 1a**), to refine our previous estimates of mutation rates in N2 wild-type (Meier et al. 2014, 2018). In line with our previous results, the average mutation rate was calculated to be ∼0.9 mutations in the diploid genome per *C. elegans* generation, which corresponds to ∼2.85 × 10^−10^ per base pair and germ cell division (Methods). Mutations were equally distributed across the genome with no evidence of mutation clustering (**Fig. 1b**). The most frequent mutations were single base insertions and deletions in homopolymeric sequences (**Fig. 1c**), indicative of replication slippage as a source of mutations in wild-type.

Investigating mutation rates across wild-type and 61 *C. elegans* DNA repair mutant backgrounds, we found that mutation rates varied depending on mutation type, making a comparison of the overall rates misleading. We thus calculated mutation rates after stratifying mutations into 1) single and multi-nucleotide variants, 2) indels smaller than 400 base pairs and 3) structural variants (SVs) (**Fig. 2, Fig. S1**). The median mutation rate across all DNA repair defective strains was close to that observed in wild-type: 0.82 heterozygous base substitutions per generation compared to 0.57 (standard deviation SD=0.04) in wild-type, 0.25 indels (0.26 (SD=0.03) in wild-type) and 0.03 SVs (0.02 (SD=0.01) in wild-type). However, 69% (42 out of 61, false discovery rate FDR=5%) of DNA repair deficient strains displayed mutation rates significantly different from wild-type in at least one mutation class, with a change of over 2-fold occurring in 44% (27 out of 61) of all lines.

**Fig. 2.**
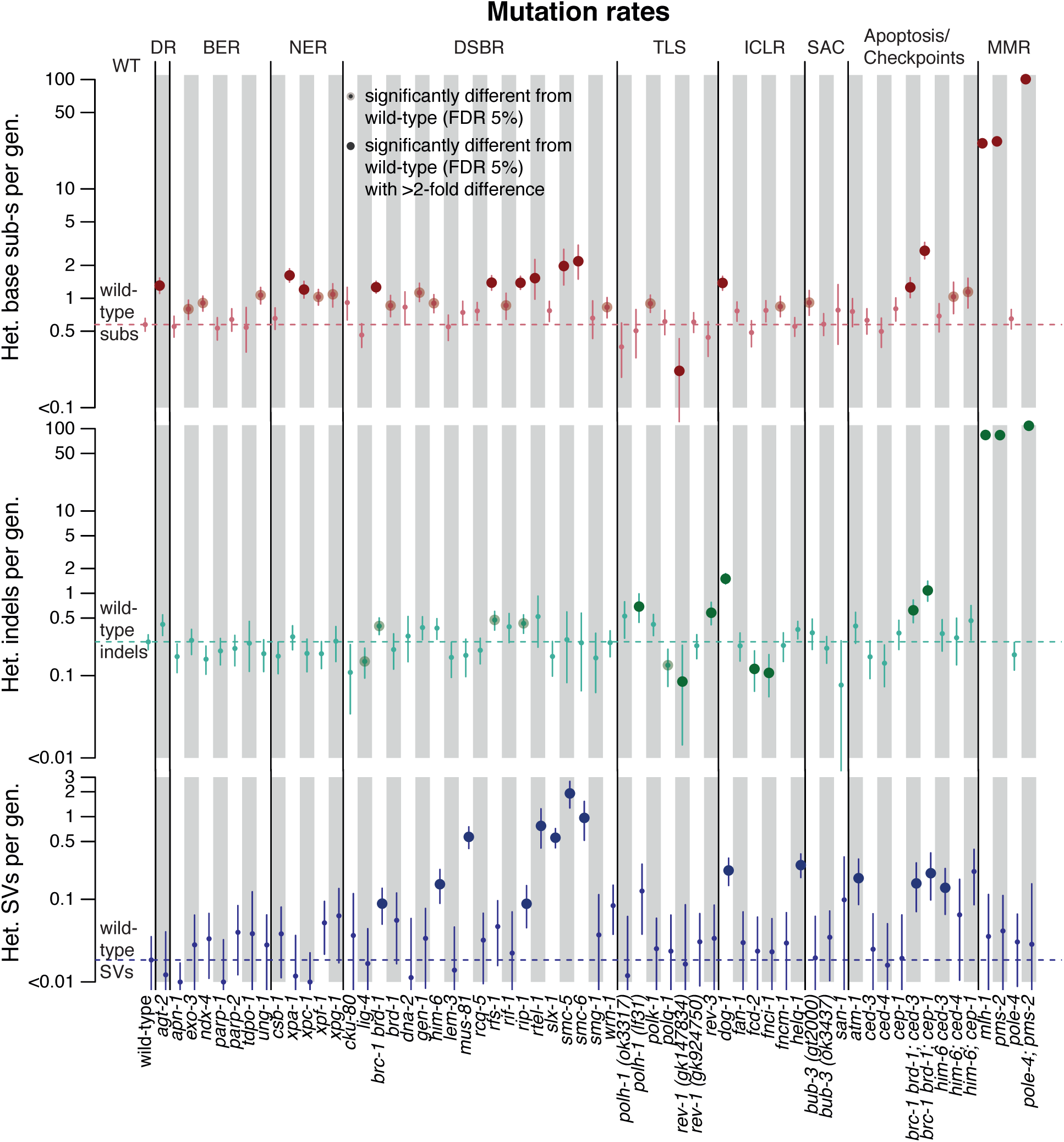
Mutation rates across 60 *C. elegans* genotypes stratified by mutation type: base substitutions, indels, and structural variants. Mutation rates are shown as number of heterozygous mutations per generation for N2 wild-type (WT), and mutants used in this study grouped by the major DNA repair pathway they contribute to; direct damage reversal (DR), base excision repair (BER), nucleotide excision repair (NER), crosslink repair (ICLR), DNA double-strand break repair (DSBR), spindle assembly checkpoint (SAC), apoptosis, and mismatch repair (MMR). Base substitutions are shown in red, indels in green and structural variants in purple. Dotted lines denote the mutation rates for wild-type. Error bars show the 95% confidence intervals; large dots represent variants with 2-fold increased or decreased mutation rates over N2 wild-type which are statistically significant with a false discovery rate (FDR) below 5%.

The strongest increase was observed in DNA mismatch repair (MMR) mutants (*pms-2* and *mlh-1*), with 25-30 times more base substitutions and a ∼100-fold increase in indels per generation (Meier et al. 2018; Zou et al. 2018; Drost et al. 2017), and even higher numbers of base substitutions in double mutants of MMR deficiency with *pole-4* (*pole-4; pms-2*) as described previously (**Fig. 2**) (Meier et al. 2018). Indels mostly comprised 1-2 base insertions or deletions in repeat sequences consistent with their etiology linked to unrepaired replication polymerase errors and slippage events (Meier et al. 2018). A ∼2 fold increase of single nucleotide variants was observed in a number of mutants defective for NER, HR, direct damage reversal (DR), and helicases, while a ∼3-5 fold increase occurred in *smc-5* and *smc-6* HR defective mutants, as well as in the *brc-1 brd-1; cep-1* triple mutant defective for the *C. elegans* orthologs of BRCA1, its binding partner BARD1, and p53 (**Fig. 2, red dots**). DNA interstrand crosslink repair deficient *dog-1/FANCJ* mutants exhibited ∼6 times more indels compared to wild-type, and several HR and TLS deficient strains showed ∼2 fold increased indel rates (**Fig. 2, green dots**). Finally, the numbers of structural variants (SVs) tended to be elevated in lines compromised for HR and for various DNA helicases (*dog-1, helq-1, him-6/BLM, rtel-1*) (**Fig. 2, blue dots**). A number of DNA repair mutants, such as *fcd-2/FANCD2* and *fnci-1/FANCI* ICL repair, as well as *lig-4* end-joining defective mutants, exhibited reduced indel rates. However, given that mutation rates for indels and SVs were already low in wild-type, we caution that the observed reductions are likely to be false discoveries. Moreover, we analysed 4-8 lines per DNA repair mutant genotype compared to 30 wild-type lines, thus the sample variance of some genotypes, especially those with mutation rates close to wild-type, may be underestimated.

### Direct damage reversal (DR), base excision repair (BER), nucleotide excision repair (NER), and single-strand break (SSB) repair

We next wished to systematically characterise the signatures and features of mutations accumulated over generations across all DNA repair pathway mutants. Mutants deficient in DR, BER, and NER did not show large changes in mutation spectra, but several small differences in particular mutation types (**Fig. 3a, Fig. S2a**).

**Fig. 3.**
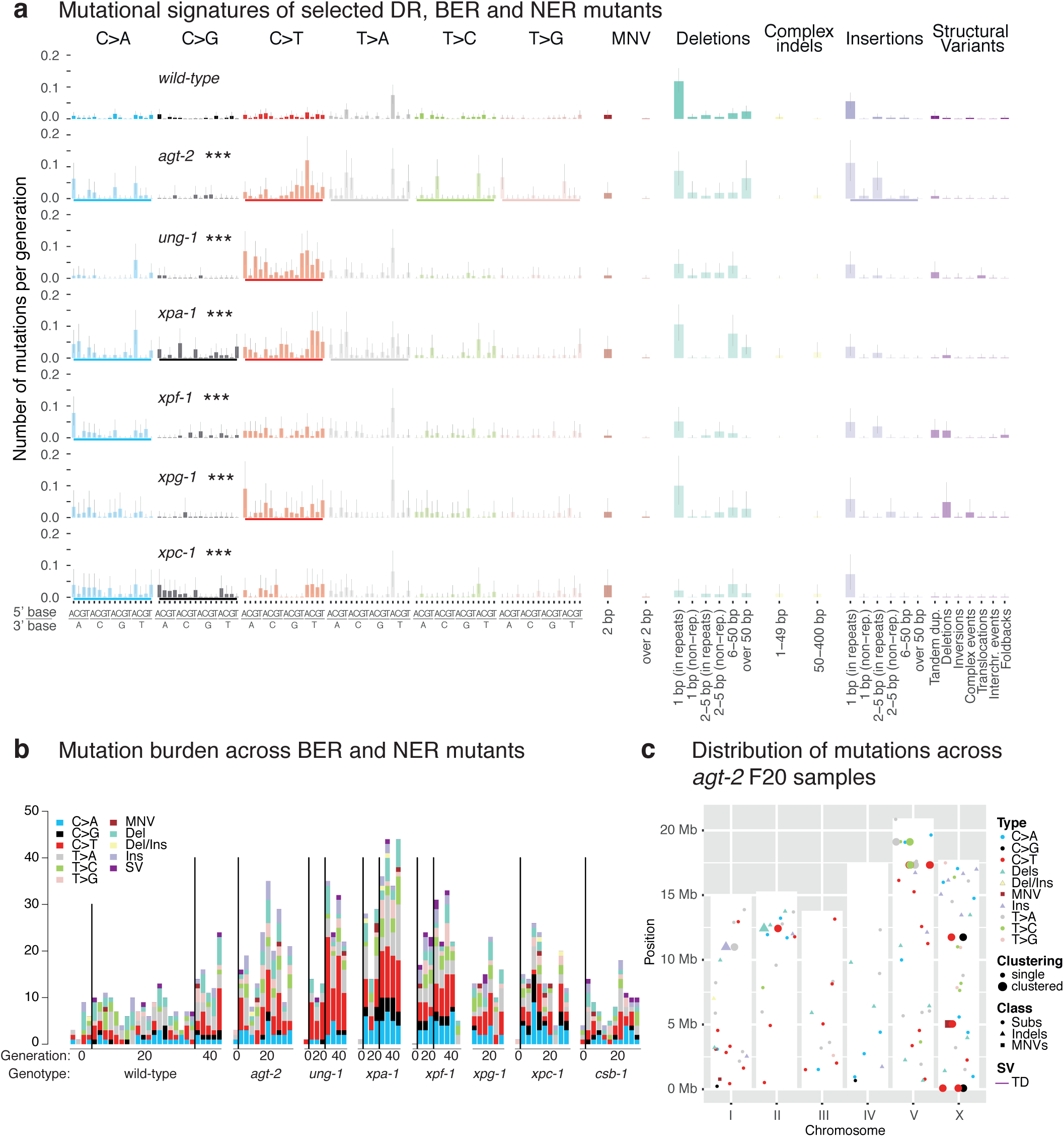
Mutation accumulation and clustering in selected *C. elegans* BER, NER, and DR mutants. **a**. Mutational signatures of BER, NER, and DR mutants that display statistically significantly different mutation spectra than wild-type shown as the number of mutations per generation across all mutation classes. Bold coloured bars below each mutation profile indicate mutation types that are different from wild-type, three stars indicate genotypes which have significantly different rates of substitutions, indels or SVs compared to those in wild-type (FDR < 5%). **b**. Total number of mutations of all classes shown for each individual sequenced line of the indicated genotype and generation. **c**. Mutation types of all classes and their location on the 6 *C. elegans* chromosomes (I-V and X) observed across all *agt-2* mutant lines. The height of the white bars corresponds to the length of each individual chromosome. Single nucleotide variants (Subs) are indicated by a dot, dinucleotide variants (DNVs) by a square, indels divided in deletions (D), insertions (I), and deletions with insertions (DI) by a triangle, and structural variants (SVs) by a line. Clustering of mutations within a single line is depicted by enlarged bold symbols.

In addition to AGT-1, which facilitates direct damage reversal by removing methyl moieties from *O*^*6*^-methyl guanine, AGT-2 encodes for a further predicted *C. elegans O*^*6*^-alkylguanine DNA alkyltransferase (Kanugula and Pegg 2001). We found a 2-fold increase in mutation rate in *agt-2* deficient lines (**Fig. 2**), owing to an elevated frequency of C>T changes caused by the mispairing of *O*^*6*^-methyl guanine with T (**Fig. 3a,b**). Interestingly, *agt-2*, exhibited a moderate degree of mutation clustering, evidenced by 7 cases of 2-3 mutations located in closer proximity to each other than expected by chance, scattered across 10 *agt-2* mutant lines (**Fig. 3c, Fig. S2b**). AGT-2 did not affect mutagenesis following methyl methanesulfonate treatment (Volkova et al. 2020). A recent study suggested a role of AGT-2 in the processing of meiotic DSBs (Serpe et al. 2019), which may explain the clustered nature of AGT-2 related mutations. We also confirmed an increase in C>T changes in mutants defective for *ung-1*, an Uracil-DNA glycosylase that excises uracil during BER (Nakamura et al. 2008; Meier et al. 2014) (**Fig. 3a,b**). Uracil is introduced via UTP mis-incorporation or cytosine deamination and pairs with adenine, which leads to C>T mutations. Other BER mutants, including mutants deficient in PARP-1 and PARP-2, the two *C. elegans* poly-ADP ribose polymerases needed for SSB repair, did not show altered mutation rates compared to wild-type (**Fig. 2, Fig. S2a**).

The NER pathway is involved in the repair of bulky DNA adducts and DNA crosslinks, both of which cause a structural distortion of the DNA double helix (Lans and Vermeulen 2011). *xpa-1, xpf-1*, and *xpg-1* lines compromised for all NER and *xpc-1* lines solely defective for global genome NER (but not *csb-1* lines uniquely defective for transcription coupled NER) showed increased mutation rates without overt changes in mutational signatures (**Fig. 2, 3a,b**; **Fig. S3a**). We speculate that this increased mutagenesis might be caused by cyclopurines induced by reactive oxygen species, and/or from exposure to ambient and fluorescent light. Comparison between the *C. elegans* NER signature adjusted for the human nucleotide composition (for methods see (Meier et al. 2018)) and COSMIC signatures SBS8, suggested to be associated with NER deficiency based on human organoid experiments (Jager et al. 2019), and SBS5 found to be associated with NER defects in urothelial cancers (Kim et al. 2016) showed no conformity (cosine similarity scores 0.64 and 0.56, respectively) (**Fig. S3b**).

### DSB repair by nonhomologous DNA end-joining (NHEJ), microhomology mediated end-joining (MMEJ) and homologous recombination (HR)

DSB repair is facilitated by several redundant pathways. HR is considered largely error-free, restoring genetic information using an intact homologous DNA strand as a repair template. End-joining pathways, classical NHEJ (c-NHEJ) and MMEJ, are typically error-prone and join free DNA ends. c-NHEJ largely acts on blunt DNA ends, while MMEJ requires short DNA resection to generate complementary 2-20 base single-stranded DNA termini to join broken DNA ends (Seol et al. 2018).

Inactivation of the core components of NHEJ, *cku-80* and *lig-4*, and of the core MMEJ component *polq-1*, did not produce significant and reliable changes in mutation rates compared to wild-type (**Fig. 2, Fig. S4a**). To study the effect of defective HR, we investigated mutation accumulation in *brc-1 brd-1* double mutants. BRC-1/Brca1 and BRD-1/Bard1 proteins function in a complex and the corresponding deficiencies are considered epistatic (Boulton et al. 2004). We report on the double mutant as our genome sequencing analysis revealed that the *brc-1* mutant strain we used also contained a *brd-1* deletion (**Table S1**). In contrast to end-joining mutants, the *brc-1 brd-1* double mutant showed increased numbers of single nucleotide variants (**Fig. 2**), small deletions between 5 and 50 bases (**Fig. 4a**), and tandem duplications (TDs) between 1.6 and 500 kbps, with a median of ∼12 kbps (**Fig. 4b,c, Fig. S5**). Overall, the mutational signature of *C. elegans* BRC-1 BRD-1 deficiency agrees with the flat profile of increased base substitutions described in HR deficient human cancers (Polak et al. 2017; Riaz et al. 2017), BRCA1 negative human lymphoblastic MA lines (Póti et al. 2019), and also resembles the pattern of SVs associated with BRCA1 loss in breast and ovarian cancers (Nik-Zainal et al. 2016; Macintyre et al. 2018; Viari et al. 2017).

**Fig. 4.**
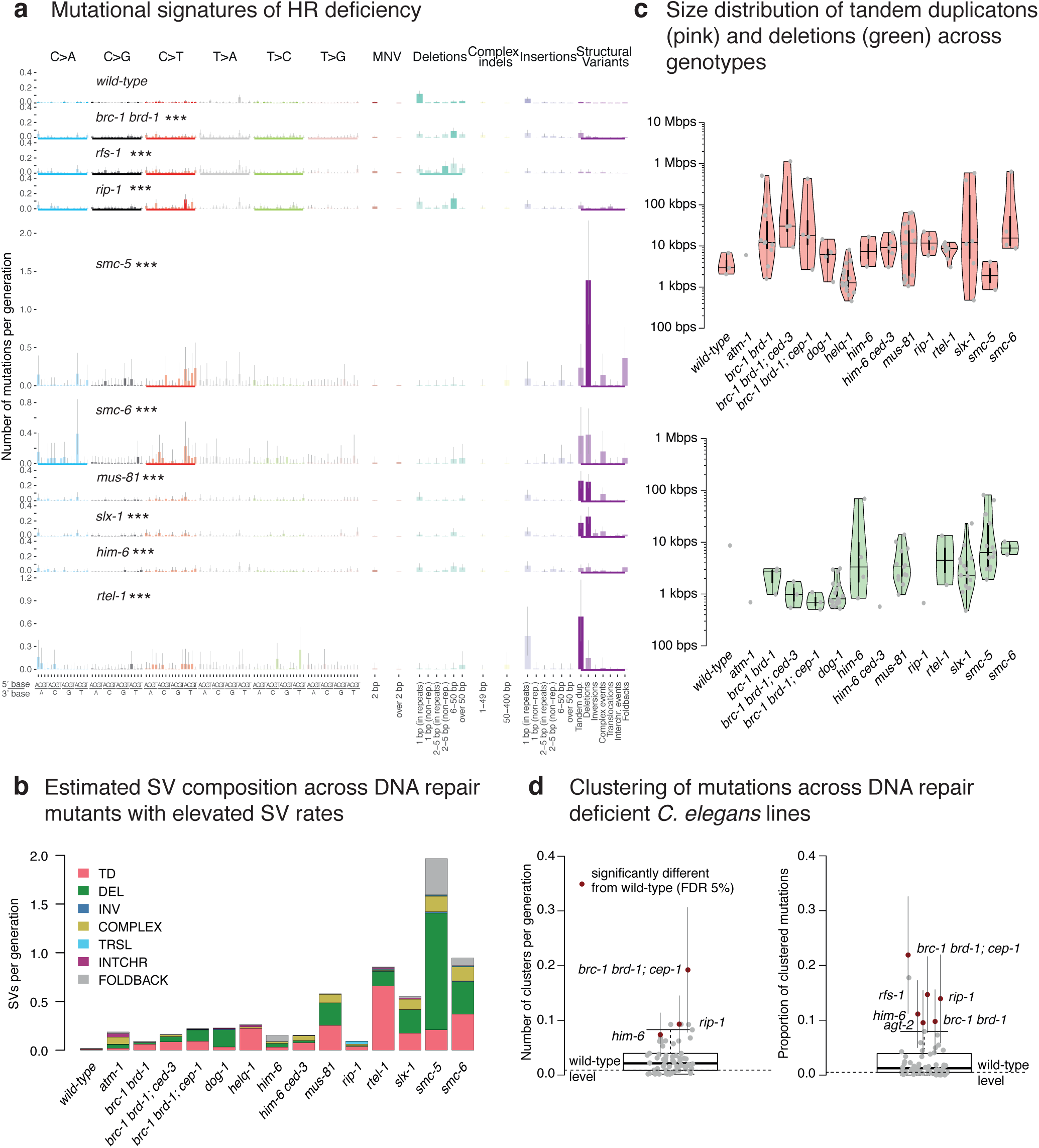
Mutational signatures and genomic features of mutations in DSBR deficient *C. elegans* lines. **a**. Mutational signatures of DSBR mutants that exhibited statistically significant differences to wild-type mutation rates displayed in numbers of mutations per generation. Bold coloured bars below each mutation profile indicate mutation types that are different from wild-type, three stars indicate genotypes which have significantly different rates of substitutions, indels or SVs compared to those in wild-type with (FDR < 5%). **b**. Estimated composition of structural variants per generation as estimated for the wild-type and genotypes with elevated SV rates. **c**. Size distributions of tandem duplications (left, pink) and deletions (right, green) across wild-type and genotypes with elevated SV rates. **d**. Clustering of mutations across DNA repair deficient *C. elegans* lines. Dots reflect the number of clusters per generation (left) or the proportion of clustered mutations. Error bars denote 95% confidence intervals. Genotypes which are significantly different from the wild-type levels (dotted black line) are shown in red.

Homologous recombination repair requires substantial end processing at the site of DSBs, which is performed by a series of nucleases creating single-stranded DNA overhangs. As mutants of *C. elegans* nucleases *rad-51, mre-11*, and *com-1* are sterile due to defects in meiotic recombination thus precluding MA experiments, we analysed mutation rates in strains deficient for the *rad-51* paralog *rfs-1* and for *rip-1* which encodes a RFS-1 interacting protein (Taylor et al. 2015). The RFS-1 RIP-1 complex is thought to stimulate the remodelling of presynaptic RAD-51-coated DNA filaments to facilitate strand invasion for recombinational repair (Taylor et al. 2015). We observed an overall 2-fold elevated mutagenesis in *rfs-1* and *rip-1* mutants (**Fig. 2**), defined by increased numbers of base substitutions in both strains, increased numbers of small deletions in *rfs-1* and increased numbers of SVs in *rip-1* (**Fig. 4a,b, Fig. S6**). Intriguingly, we observed three ‘translocation type’ events in *rip-*1 but not in any other MA line we analysed (**Fig. 4b, Fig. S6, Fig. S7**). We deduced that these events involved templated insertions of 200-4000 bp sequences, which showed strong homology to multiple genomic regions, to one such homeologous region *in cis* located as far as 275 kb away from the donor sequence on the same chromosome, accompanied by a deletion of several hundred basepairs at the homeologous acceptor site (**Fig. S7**). These templated insertions may be explained by strand invasion into homeologous template DNA, in line with the pro-recombinogenic role of RIP-1 in mediating RAD-51 dissociation from invading strands (Taylor et al. 2015).

SMC-5 and SMC-6 are components of a ring shaped cohesin complex considered to tether broken DNA strands to the repair template on the sister chromatid to facilitate HR (Bickel et al. 2010). *smc-5* and *smc-6* mutants, which have been shown to exhibit defects in meiotic recombination between sister chromatids (Bickel et al. 2010; Hong et al. 2016) showed an increased rate of base substitutions and SVs, largely comprised of deletions, tandem duplications, and complex rearrangements (**Fig. 2, 4a,b, Fig. S6)**. In agreement with a high preponderance of SVs, these lines could only be propagated for up to 5 generations before succumbing to sterility. In contrast to *brc-1 brd-1* mutants, *smc-5* and *smc-6* mutants exhibited a high proportion of large ∼10 kb deletions (**Fig. 4c)**. Given the role of the ring-shaped SMC-5-6 complex in enforcing close proximity of damaged and template DNA, we hypothesize that double-strand break induced HR is initiated in the absence of the SMC-5-6 complex, but the displacement loop (D-loop) formed following strand invasion falls apart prematurely resulting in the loss of genetic material.

The structure-specific nucleases MUS81 and SLX1 act in conjunction to process Holliday Junctions, key four-way DNA intermediates of HR (Agostinho et al. 2013; Wyatt et al. 2017, 2013). *slx-1* and *mus-81* mutants displayed similar mutational signatures characterized by increased numbers of base substitutions and SVs, with large deletions and TDs being most prevalent (**Fig. 2, 4a,b, Fig. S8**). In contrast, the absence of GEN-1, a canonical Holliday Junction resolvase (Bailly et al. 2010), or LEM-3, the ortholog of mammalian Ankle1, recently implicated in the processing of recombination intermediates that persist beyond anaphase (Hong et al. 2018), did not yield overt changes in mutation rates or signatures **(Fig. 2, Fig. S4a)**.

DNA helicases are enzymes that unwind double-stranded DNA. Among their multiple roles in HR, they contribute to the unwinding of D-loop structures, a function especially important when a broken DNA end invades a template strand with imperfect sequence homology, thus preventing recombination with homeologous sequences. To investigate mutation patterns induced by helicase deficiencies, we analysed mutants defective for the three *C. elegans* RecQ helicases: *him-6* - the ortholog of the mammalian Bloom syndrome gene which encodes for a helicase involved in HJ resolution and prevention of crossover recombination (De Muyt et al. 2012; Wicky et al. 2004); *wrn-1*, the ortholog of Werner’s syndrome gene which encodes for a helicase possessing an N-terminal 3’-5’ exonuclease domain and capable of resolving aberrant DNA structures with 3’ recessed ends (Ozgenc and Loeb 2005; Lee et al. 2004); and *rcq-5*, the ortholog of human RECQ5, which displaces RAD-51 from single-stranded DNA and thus prevents excessive recombination (Paliwal et al. 2014). In addition, we analysed lines deficient for *rtel-1* which encodes a conserved helicase involved in genome stability and telomere maintenance (Uringa et al. 2011). While *rcq-5* and *wrn-1* mutants did not show increased mutagenesis, *him-6* mutants demonstrated 8-fold elevated SV rates compared to wild-type, with 0.15 SVs per generation (SD = 0.03) (**Fig. 2, 4a, b, Fig. S4a, Fig. S9**). Even more SVs were observed in *rtel-1* mutants with an estimated rate of 0.8 TDs per generation (SD = 0.2) (**Fig. 2, 4a,b**), which spanned on average 8 kbps and were generally smaller than TDs observed in *brc-1 brd-1* mutants (**Fig. 4c**). In addition, *rtel-1* deficiency led to ∼2.5 fold increase in base substitutions (**Fig. 2, 4a**). RTEL-1 has a unique role in preventing heterologous recombination during break-induced repair and in promoting non-crossover products (León-Ortiz et al. 2018; Youds et al. 2010). Interestingly, loss of mammalian RTEL1 yields a high number of large deletions and complex rearrangements as a result of excessive crossover and heterologous recombination (León-Ortiz et al. 2018), unlike our data which showed a more simple, tandem duplication signature (**Fig. 4, Fig. S8**). In our experiments, *C. elegans rtel-1* mutants did not grow beyond F15, and most lines became sterile within 5 generations (F5) (**Table S1**), suggesting that the absence of RTEL-1 may lead to accumulation of SVs incompatible with organismal viability.

Investigating the genomic context of structural variants, we did not observe any overt changes in the presence of microhomology at the breakpoints compared to wild-type (**Fig. S4b,c**). In addition, we confirmed that SVs across almost all HR deficient genetic backgrounds tended to be associated with repetitive DNA regions (**Fig. S4d**) in line with previous reports (Puddu et al. 2019).

Among the HR mutants, we note that *brc-1 brd-1, rfs-1, rip-1, smc-5, and smc-6* display elevated levels of base substitutions. Increased base substitutions were also observed in *BRCA1* defective human lymphoblastic MA lines (Póti et al. 2019). Moreover, *brc-1 brd-1, him-6, rip-1*, and *rfs-1* exhibit evidence of mutational clustering (**Fig. 4d**) with 10% -15% of base substitutions occurring within distances smaller than expected by chance (**Methods**). Clusters of mutations may arise through error-prone polymerases reading across lesions (Supek and Lehner 2017; Stone et al. 2012; Roberts et al. 2012). In addition, the NHEJ or MMEJ error-prone DSB repair pathways can also generate clustered mutations when DNA ends are joined together (Chen et al. 2012).

### Translesion synthesis (TLS)

TLS polymerases are specialised DNA polymerases able to move across and insert nucleotides opposite damaged bases. Depending on the inserted nucleotide, this leads to error-free or error-prone lesion bypass (Sale 2013). *C. elegans rev-3/REV3L* mutants, deficient in the catalytic subunit of polymerase ζ, accumulated increased numbers of 50-400 bp deletions (**Fig. 2, 5a, Fig. S10**). Similarly, *polh-1*/*POLH (lf31)* and *polh-1/POLH (ok3317)*, DNA polymerase η mutants, displayed 50-400 bp deletions, with only *polh-1(lf31)* reaching clear statistical significance over the generations tested (**Fig. 2, Fig. S10**). Our data suggest that REV-3 and likely POLH-1 prevent DNA breaks by reading across damaged bases that seemingly also occur in the absence of exogenous DNA damage which triggers TLS activity and produces a similar signature. Our results on REV-3 and POLH-1 are in line with earlier findings in *C. elegans* (Schimmel et al. 2019; Roerink et al. 2014; van Schendel et al. 2016; van Bostelen et al. 2020) and studies in yeast and mammalian cells (Lawrence and Hinkle 1996; Lange et al. 2012).

**Fig. 5.**
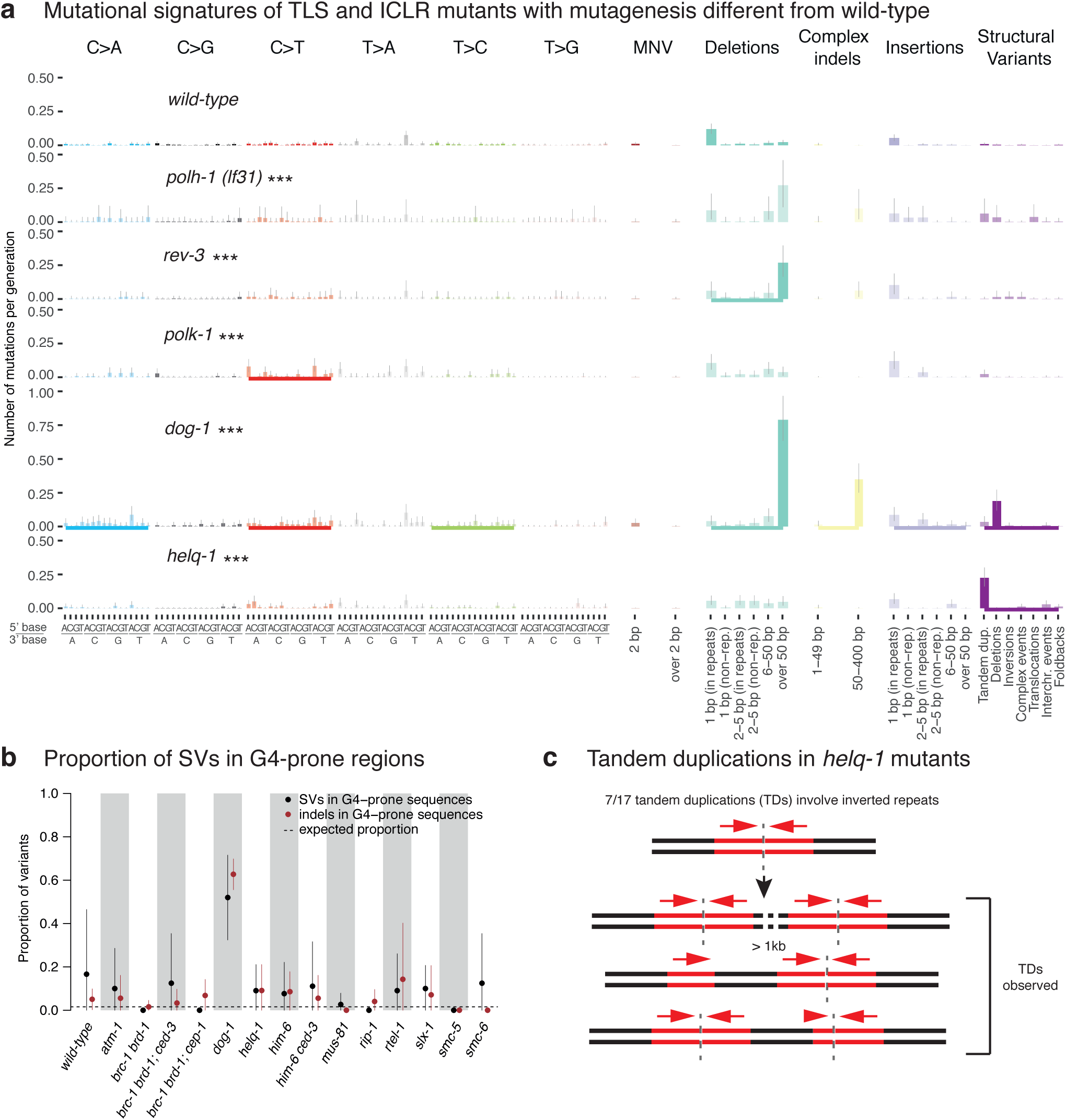
Signatures and genomic features of mutations in TLS and ICLR deficient *C. elegans* lines. **a**. Mutational signatures of TLS and ICLR mutants that exhibited statistically significant differences to wild-type mutation rates displayed in numbers of mutations per generation. Bold coloured bars below individual mutation profiles indicate mutation types that are different from wild-type, three stars indicate genotypes which have statistically significantly different rates (FDR < 5%) of substitutions, indels or SVs compared to those in wild-type. **b**. Proportion of indels (brown) and SVs (black) in G-rich regions in wild-type and across genotypes with elevated rates of SVs. Dotted line represents the proportion of variants falling into these regions as expected by chance. **c**. Tandem duplications in *helq-1* mutants.

In addition, we observed a slightly increased base substitutions rate, namely for C>T changes, in *polk-1*, which may indicate a role in error-free bypass of endogenously induced guanine modifications (Choi et al. 2006) (**Fig. 5a, Fig. S10**). Mutants defective in REV-1 TLS polymerase, as well as *polq-1/POLQ* polymerase, did not demonstrate a significant and reliable change in mutagenesis compared to wild-type (**Fig. S10**).

### Mutation accumulation in mutants deficient for DNA crosslink repair

The repair of DNA interstrand crosslinks (ICL) provides a formidable task. It involves the Fanconi Anaemia (FA) proteins required for sensing ICLs and assembling various repair factors at the site of damage (Kottemann and Smogorzewska 2013; Hashimoto et al. 2016; Ceccaldi et al. 2016). Here we investigate mutant lines defective for *fncm-1*/FANCM, a helicase involved in DNA damage recognition, FANCI-1/FANCI, and FCD-2/FANCD2, the key Fanconi repair proteins ubiquitinated by an E3 ubiquitin ligase complex and thought to assemble proteins required for ICL processing. We did not observe overt differences in mutagenesis between *fcd-2, fncm-1*, or *fnci-1* mutants and wild-type, suggesting that the *C. elegans* Fanconi Anemia ICL repair pathway does not significantly contribute to the repair of DNA damage that occurs under normal, unchallenged growth conditions (**Fig. 2, Fig. S11a**). FAN-1/FAN1 is a conserved structure-specific DNA nuclease that can resolve ICLs independently of the FA pathway (Jin and Cho 2017; MacKay et al. 2010; Kratz et al. 2010). As for the core FA components, we did not find overt mutational changes in *fan-1* mutant lines (**Fig. 2, Fig. S11a**).

DOG-1, the *C. elegans* ortholog of the mammalian FANCJ helicase, facilitates error-free replication through DNA tertiary structures formed by G-rich DNA sequences referred to as G-quadruplexes (Cheung et al. 2002; Youds et al. 2008; Tarailo-Graovac et al. 2015). Indeed, *dog-1* mutants exhibited increased mutagenesis (**Fig. 2**) with 6-fold higher numbers of 50-400 base pair indels and 13 fold more SVs, predominantly deletions (**Fig. 5a, Fig. S9**). Across all 11 *dog-1* deficient samples, 81% of long deletions (17/21) and 78% (109/139) of shorter, 50-400 bp deletions overlapped with one of the 4291 regions in the *C. elegans* genome predicted to form G-quadruplex structures (Marsico et al. 2019), in line with previous reports (Tarailo-Graovac et al. 2015) (**Fig. 5b, Fig. S9, S11B**). We found that G-quadruplex induced deletions occurred at a frequency of about 1 lesion per generation in *dog-1* mutants. We rarely observed SVs in regions containing G-quadruplex forming sequences in any other DNA repair mutants (**Fig. 5b, Figs. S5, S6, S8, S9**), including *him-6*, encoding the *C. elegans* ortholog of the mammalian BLM helicase (**Fig. S9**), which has been shown to prevent replication fork stalling at G-quadruplex sites in human and murine cells in conjunction with FANCJ (van Wietmarschen et al. 2018).

Another helicase mutant that displayed a distinct phenotype was *helq-1*/HELQ, encoding for a conserved helicase and thought to act in DNA crosslink repair in a pathway separate from FCD-2 (Muzzini et al. 2008). In addition, *helq-1* has been shown to be synthetic lethal with *rfs-1* due to its role in resolving DSB repair intermediates during meiosis (Ward et al. 2010). *helq-1* deficient lines showed an increased proportion of tandem duplications (TDs) compared to most other strains (**Fig. 4b, Fig. 5a, Fig. S5**). TDs ranged in size between 457 and 8089 bp with a median of 1270 bp (**Fig. 4c**). The mutational spectrum of *helq-1* differed from that of *brc-1* mutants, in which deletions of 6-50 bp and TD (with a median size of ∼12 kbps) were observed with comparable frequency (**Fig. 4b,c, Fig. S5**). Interestingly, 41% (7/17) of TDs in *helq-1* mutants were associated with inverted repeat sequences (**Table S2**). To investigate how TDs present at inverted repeats relate to DNA replication directionality, we used the closest origin of replication as a reference point and tested the correlation between the orientation of the TD breakpoints and the direction of leading strand synthesis. Out of the 7 inverted repeat-associated TDs we could determine the directionality of replication in 5 cases (**Methods, Table S2**). In 4 cases TD oriented in line with leading strand replication. In 3 of these inverted repeats were present downstream of the TD, in 1 case upstream (**Methods, Table S2**).

At present, we can only speculate how these tandem duplications might arise. It is likely that the genesis of TDs involves microhomology-mediated break-induced replication. Inverted repeats may form secondary stem-loop structures that impede replication fork progression (**Fig. 5c, Fig. S12**). In the case of the presence of a stem-loop structure in the DNA ahead of a replication fork, stalling could occur during the attempt to unwind the secondary DNA structure. The replication machinery may re-initiate at a downstream template, resulting in a duplicated region, before successfully replicating through the stem loop structure in a second attempt. In such a scenario, tandem duplications would always occur upstream of the inverted repeat (**Fig. 5c, Table S2**). Alternatively, an inverted repeat could more readily adopt a stem loop structure in the extended single-stranded region of the lagging strand, which is particularly prone to form secondary structures (**Fig. 5c, Fig. S12**). Such stem loop structures would be prime substrates for HELQ which has been shown to bind to single-stranded DNA and act as a 3’ to 5’ helicase thus facilitating the dissolution of the stem loop (Marini and Wood 2002). In the absence of HELQ activity, these stem loops could be recognised by a nuclease (**Fig. S12**) and a resulting single-strand break might facilitate the invasion into the freshly replicated leading strand DNA and prime break-induced replication. Recapturing the original template (**Fig. S12**, step 3) after break-induced replication would restore the fork, resulting in a tandem duplication in only one of the two chromatids (**Fig. S12**). Crucially, the position of the nucleolytic cut, upstream, downstream or within the inverted repeat and the position at which invasion into the template strand occurs would determine the breakpoint of the tandem duplication.

In summary, our data suggest that the DNA helicases HELQ-1 and DOG-1 are required to facilitate replication fork passage through distinct secondary structures (**Fig. 5b,c**). While DOG-1 is needed for the passage through G-rich structures (Cheung et al. 2002), HELQ-1 may help to overcome stem loop structures, possibly on the lagging strand.

### Mutations and subtelomeric chromosome fusions in ATM-1 defective strains

ATM is a conserved PI3 kinase involved in DNA damage checkpoint regulation and telomere homeostasis. ATM deficiency has been reported to be associated with shorter telomeres from yeast to mammalian cells. Moreover, in ATM deficient yeasts, *C. elegans* and *Drosophila*, the latter depending on retrotransposon transposition rather than telomerase activity for telomere maintenance, chromosome fusions have been observed cytologically or through sequencing of PCR products across chromosomes (Chan and Blackburn 2003; Mieczkowski et al. 2003; Bi et al. 2004; Silva et al. 2004; Oikemus et al. 2004; Lee et al. 2015). *C. elegans atm-1/ATM* mutants are hypersensitive to ionizing radiation (IR) (Jones et al. 2012; Boulton et al. 2002). In addition, *atm-1* lines propagated over multiple generations have been described to display a stochastic *him* (<u>h</u>igh incidence of <u>m</u>ales) phenotype, an indicator of meiotic chromosome mis-segregation associated with sex chromosome to autosome fusions (Jones et al. 2012; Ahmed and Hodgkin 2000). Investigating mutation rates in *atm-1* lines grown for 20 generations, we observed 2-fold elevated numbers of SVs, predominantly inversions (**Fig. 2, Fig. 6a**). This increased incidence of SVs agrees with previous estimates of mutation rates in *atm-1* mutants which were based on scoring the number of essential mutations in *atm-1* backgrounds (Jones et al. 2012). Interestingly, 4 of the 5 *atm-1* lines grown for 20 generations carried SVs (with over 70% (9/11) of all observed SVs) localised in subtelomeric regions **(Fig. 6b, Fig. S5)**, unlike any other DNA repair deficient mutants examined. Most subtelomeric SVs (8/9) could be classified as complex rearrangements with at least 2 overlapping or adjacent events, often associated with copy number changes **(Fig. 6c, Fig. S5, Fig. S13)**. Moreover, 4/5 complex rearrangements involving a single chromosome end displayed a loss of telomere sequences, suggesting a possible end-to-end fusion between homologous chromosomes. Furthermore, we observed 2 cases of interchromosomal rearrangements between different autosomes with breakpoints in subtelomeric regions associated with deletion of telomere sequences and copy number alterations **(Fig 6b, Fig. S5, Fig. S13)**. None of the *atm-1* lines carried translocations or amplifications of the genomic regions termed TALT1 or TALT2, amplified in survivors of telomerase-deficient *C. elegans* strains and considered to be utilized as templates for an alternative (telomerase independent) telomere lengthening (ALT) mechanism (Seo et al. 2015). Similarly, we did not observe translocation events associated with *atm-1* SVs **(Fig. S5, Fig. S13)** making templated telomere maintenance from interstitial telomere sequences buried in the genome unlikely.

**Fig. 6.**
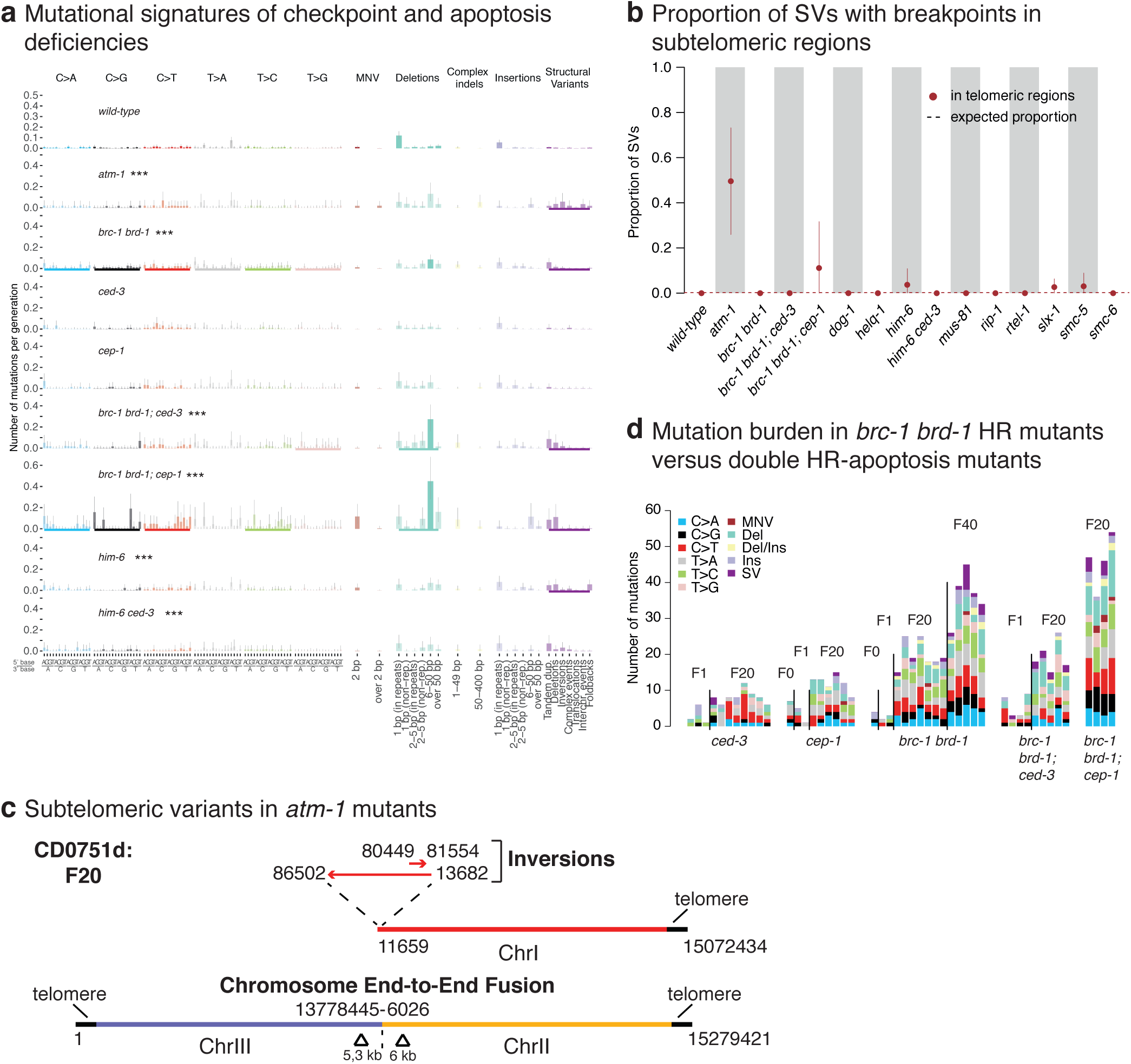
Signatures and genomic features of mutations in DNA damage checkpoint and apoptosis deficient *C. elegans* lines. **a**. Mutational signatures of mutants that exhibited statistically significant differences to the wild-type mutation rates. Bold coloured bars below each mutation profile indicate mutation types that are different from wild-type, three stars indicate genotypes which have significantly different rates of substitutions, indels or SVs compared to those in wild-type with (FDR < 5%). **b**. Proportion of SVs with breakpoints falling into subtelomeric regions across wild-type and genotypes with elevated SV rates. Dotted lines represent the fraction of variants expected to hit subtelomeric regions by chance. **c**. Examples of structural variants in *atm-1* mutants. **d**. Quantification of mutation burden of the indicated lines after 20 or 40 generations as indicated.

We speculate that loss of *C. elegans atm-1* function could lead to reduced recruitment or access of telomerase to the shortest telomeres consistent with studies in tel1/ATM mutant yeast (Hector et al. 2007; Bianchi and Shore 2007; Ritchie and Petes 2000), thereby leading to telomere loss and chromosome fusions. Access of telomerase to its telomeric substrate could be inhibited by *atm-1* loss due to reduced end resection, a mechanism also discussed for *atm*^*-/-*^ dependent telomere shortening and fusions in mammalian cells (Wu et al. 2007).

### p53 and apoptosis defective strains

In the presence of excessive DNA damage, cells can trigger the p53 pathway to activate their apoptotic demise. CED-3 and CED-4, orthologs of a human caspase and the APAF1 protein, are essential for DNA damage induced and developmental apoptosis (Gartner et al. 2000). In contrast, CEP-1, the *C. elegans* p53 ortholog is specifically required for DNA damage induced apoptosis (Schumacher et al. 2001; Derry et al. 2001) (**Fig. S14**). We did not observe increased mutagenesis in *ced-3, ced-4*, and *cep-1* defective mutants indicating no major role of DNA damage induced apoptosis for preventing mutagenesis under unchallenged growth conditions. Our dataset also includes deletions of spindle assembly checkpoint (SAC) genes. The spindle assembly checkpoint delays anaphase progression until all chromosomes are correctly attached to the mitotic spindle apparatus thus ensuring faithful chromosome segregation (Encalada et al. 2005). In addition, SAC has been implicated in DSB repair consistent with *bub-3* and *san-1* SAC mutants exhibiting IR sensitivity (Bertolini et al. 2017). Lines deficient for *bub-3* and *san-1*, corresponding to human BUB3 and BUB1B proteins, respectively (Hajeri et al. 2008), did not show increased mutagenesis (**Fig. 2, Fig. S14**).

Having observed distinct mutational patterns in HR deficient mutants, we wanted to test whether combining HR mutants with apoptosis and or DNA damage response deficiency would lead to increased or altered mutagenesis. Increased *cep-1/p53* dependent germ cell apoptosis has been reported in a number of HR mutants such as *him-6* and *brc-1*, suggesting that a higher number of nuclei carry increased or unrepaired DNA damage which might be eliminated by *cep-1* dependent apoptosis *(Wicky et al. 2004; Boulton et al. 2004; Grabowski et al. 2005; Adamo et al. 2008)*. Double mutants of *him-6* with *ced-3* (6 lines), *ced-4* (3 lines) or *cep-1* (3 lines) did not display changes in mutation rates or spectra compared to *him-6* single mutants (**Fig. S14**. In contrast, 4 *cep-1; brc-1 brd-1* lines (but not the 5 *brc-1 brd-1; ced-3* lines) showed an increased rate of mutagenesis compared to *brc-1 brd-1* alone **(Fig. 6a, d)**. Specifically, *cep-1* inactivation exaggerated the mutational features of the *brc-1 brd-1* mutant, leading to increased incidence of small deletions and structural variants **(Fig. 6a)**. We also observed that clustering of base substitutions in the triple mutant was twice as prominent as in the *brc-1 brd-1* double mutant line, with 20% of mutations in clusters **(Fig. S14)**. Thus, at least for HR deficiency conferred by *brc-1 brd-1*, additional *cep-1* inactivation increases mutation clustering. Given that apoptosis deficiency in *brc-1 brd-1; ced-3* lines does not result in increased mutagenesis, we speculate that increased mutagenesis in *brc-1 brd-1; cep-1* lines might be associated with a role of CEP-1 independent of apoptosis regulation. CEP-1 could either trigger the cell-cycle checkpoint or facilitate more efficient DNA repair.

## Discussion

Here we systematically catalogued the mutational characteristics of DNA repair deficiencies across all conserved *C. elegans* DNA repair and damage response pathways in inbred lines propagated for up to 40 generations. Our data provide a comprehensive picture of the contributions of various DNA repair and damage response pathways towards genome integrity in an experimental system under spontaneous, endogenous conditions not challenged by mutagen exposure. Except for mismatch repair deficiency (Meier et al. 2018; Puddu et al. 2019), defined repair deficiencies only lead to modest effects, with a 2-5 fold increase of mutations in 44% of all strains tested, with notable examples in almost every pathway. Alkylguanine alkyltransferase, Uracil glycosylase and NER deficiency are associated with increased acquisition of base changes. TLS mutants *rev-3*(pol ζ) and *polh-1*(pol η) show elevated numbers of 50-400 bps deletions. Interestingly, HR deficiency can manifest in different ways. *brc-1/Brca1* and *rad-51* paralog mutants show elevated mutagenesis across all types of mutations Other HR mutants inactivating the MUS-81 and SLX-1 nucleases and the HIM-6/BLM, HELQ-1 and RTEL-1 helicases are associated with increased numbers of SVs. DOG-1 has a unique role preventing deletions next to G-rich sequences, while the HELQ helicase may contribute to faithful replication across secondary DNA structures such as inverted repeats. RIP-1 appears to avert templated insertion into homeologous sequences. The ATM-1 checkpoint kinase prevents chromosome end-to-end fusions. Finally, deficiency of the *p53* like gene *cep-1* exacerbates mutagenesis caused by HR defects.

### Redundancy of DNA repair pathways

It is well established that thousands of DNA lesions occur spontaneously during each cell-cycle and that the vast majority of DNA lesions are repaired. Thus, the absence of significant mutagenic effects in the majority of genotypes tested underpins a high level of redundancy among different DNA repair pathways. Moreover, it may require the combined deficiency of multiple DNA repair pathways to trigger excessive mutagenesis in the germline. Such reasoning is in line with our recent whole genome analyses showing that multiple pathways act in concert to repair DNA lesions caused by the exposure to known genotoxins, with deficiencies of different pathways potentially leading to increased mutagenesis and/or altered mutagenic signatures (Volkova et al. 2020). Equally, latent defects such as those caused by the deletion of the non-essential polymerase subunit *pole-4*, only become apparent in conjunction with a DNA repair deficiency (Meier et al. 2018). While *pole-4* alone does not cause increased mutagenesis, combining this mutant with MMR causes mutagenesis beyond what is observed upon MMR alone (Meier et al. 2018). Many cases of DNA repair pathway redundancies have been described, for instance, simultaneously defective TLS, NER and TMEJ renders *C. elegans* sensitive to normal levels of daylight (van Bostelen and Tijsterman 2017).

In contrast to a recent large-scale mutation accumulation screen in budding yeast (Puddu et al. 2019), we did not observe widespread copy number changes in our analysis, likely because most such changes are incompatible with viability and fertility of a multicellular organism such as *C. elegans*. It is possible that nematodes suffering from gross chromosomal alterations are lost during propagation across generations. However, using our experimental setup, we were previously able to detect instances of severe chromosomal rearrangements, including complex chromosome fusion events that contain scars indicative of breakage-fusion-bridge cycles and chromothripsis (Meier et al. 2014). Nevertheless, it is likely that we underestimate mutation rates, especially in strains that could not be propagated for 40 generations, namely those defective for the RTEL-1 helicase and the SMC-5 and SMC-6 cohesion proteins.

### Mechanistic insights

Our detailed characterisation of mutation rates, mutational signatures and localised mutation features combined with the known enzymology of many repair enzymes provides mechanistic insights: Our data confirm the specific role of the DOG-1/FANCJ helicase in unwinding G-quadruplex forming sequences (Kruisselbrink et al. 2008), and we show that this feature is unique among the helicase mutants we analysed. We also reveal a specific role of the HELQ-1 helicase frequently in the context of repetitive sequences such as inverted repeats. Our breakpoint analysis of tandem duplication associated with HELQ-1 deficiency, together with the known biochemical role of HELQ helicases, supports a model where this helicase might be loaded to the single stranded area of the lagging strand and help to dissolve palindromes and facilitate lagging strand synthesis. Furthermore, cases of unique gene conversion events into homeologous sequences in *rip-1* mutants support a role of RAD-51 paralogs in preventing homeologous recombination. This is in line with biochemical activity of the RFS-1 RIP-1 paralog complex in remodelling presynaptic RAD-51-containing DNA filaments to facilitate stand invasion for recombinational repair (Taylor et al. 2015). Finally, we provide evidence that the ATM-1 checkpoint kinase has a specific role in protecting sub-telomeric repeats from DSBs and preventing deletions, inversions and chromosome fusions.

Based on their mutational signatures, strains defective for homologous recombination can be broadly grouped in two classes. First, BRC-1 and RAD-51 paralog mutants show elevated numbers of point mutations, as well as increased numbers of small deletions, and structural variants. A similar pattern was observed in a study of HR knockouts in chicken DT40 cell lines (Póti et al. 2019). We suspect that increased point mutations might be a scar indicative of error prone translesion synthesis, necessary when damaged bases are neither repaired by BER and NER nor by replication fork reversal which is linked to recombinational repair (Quinet et al. 2017). Point mutations and small deletions also occur when HR is replaced by more error-prone NHEJ or MMEJ pathways, the latter being associated with the occurrence of small deletions in human BRCA1 mutants (Viari et al. 2017). Conversely, deficiencies of other HR proteins, like SLX-1 and MUS-81, and helicases including HIM-6, RTEL-1 and HELQ-1, are associated with a specific increase of SVs. It is tempting to speculate that these proteins may not have a role in HR pathways directly linked with DNA replication (see below).

### Nature of germ cell divisions and mutagenesis

It is important to keep in mind that germ cell mutagenesis might occur at different stages of the *C. elegans* life cycle. The nature of germ cell divisions is fundamentally different across various developmental stages. During the invariant embryonic development of *C. elegans*, the germ cell lineage is defined by 3 asymmetric cell divisions, and from the first zygotic cell division onwards, a single posterior daughter cell always defines the germ line, which finally splits as a part of the 4^th^ germ cell division into the two founder cells (Strome 2005). Starting from the L1 larval stage, each of these founders within a timeframe of three days expands to form one of the two gonads comprising ∼1000 germ cells (Strome 2005). Embryonic germ cell divisions occur very rapidly within a timeframe of less than 20 minutes, and cells are largely refractory to DNA damage checkpoints. Evidence exists that translesion polymerases are particularly important during this stage (Kim and Michael 2008; Roerink et al. 2012). The first cell-cycle in developing germ lines occurs after an extended period of transcriptional quiescence and synchronized transcriptional onset appears particularly challenging for genome integrity (Wong et al. 2018; Ou et al. 2019). In contrast, germ cells residing in the adult germ line are subject to cell-cycle and apoptosis checkpoints, and take an excess of 10 hours to complete (Gartner et al. 2000). It appears possible that the increased number of mutations observed in *cep-1 brc-1* double mutants reflects a role of the CEP-1 p53 like protein in preventing excessive mutagenesis. CEP-1 is expressed in mitotically dividing germ cells, as well as in late pachytene where late stages of meiotic recombination occur. Given that we did not observe comparable mutagenesis in *brc-1* and apoptosis defective double mutants, apoptosis being restricted to pachytene cells, we speculate that mutagenesis might reflect a role of CEP-1 in mitotically dividing germ cells, possibly affecting the cell-cycle checkpoint or DNA damage response. Finally, it is reasonable to suggest that many lesions we observe to accumulate in our transgenerational setup occur in haploid germ cells, especially during meiosis. Indeed, it appears plausible that many SVs might be associated with meiotic recombination. A large excess of DSBs are generated by the SPO-11 nuclease during meiosis, and typically only one DSB per chromosome pair matures into a crossover to facilitate the exchange of genetic information between maternal and paternal chromosomes (Hillers et al. 2017). Many SVs we observed in HR mutants, particularly those that did not demonstrate an excessive accumulation of point mutations, likely result from faulty recombinational events during meiosis: For instance, the SLX-1 and MUS-81 nucleases, and the HIM-6 helicase contribute to the resolution of meiotic Holliday junction intermediates (Agostinho et al. 2013; Saito et al. 2013; O’Neil et al. 2013). In contrast, BRC-1 and the SMC-5/6 complex are implicated in the repair of the excess meiotic DSBs not engaged in crossover recombination (Hong et al. 2016).

## Methods

### *C. elegans* strains, propagation and maintenance

All *C. elegans* strains used in this study (**Table S1**) were backcrossed 6 times against the wild-type N2 Bristol reference strain TG1813 (Meier et al. 2014). Strains were then grown for 20 or 40 generations as described previously (Meier et al. 2014) propagating a minimum of five lines per genotype in parallel with at least three used for sequencing. Following clonal expansion of the final generation, genomic DNA was isolated using the ChargeSwitch® gDNA Mini Tissue Kit (Invitrogen).

### Variant analysis

Illumina sequencing, variant calling, post-processing filtering, and data analysis were performed as described previously (Meier et al. 2018; Volkova et al. 2020) with WBcel235.74.dna.toplevel.fa as the reference genome (http://ftp.ensembl.org/pub/release-74/fasta/caenorhabditis_elegans/dna/).

### Mutation rates and mutational signatures calculations

Per-generation mutation rates for each genotype were estimated using additive non-negative Poisson regression using samples with generation higher than 1. Every sample *i, i* = *1*, … *N* out of *N* = *513* was assigned a number of mutations of the category of interest, *Y*_i_ ∈ *N* ∪ {*0*}. For a vector of mutation counts *Y*, 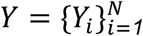, we calculated the mutation rates per generation for each genotype using the following model:

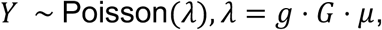

where 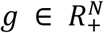 is the adjusted number of generations which takes into account the 25% chances of a heterozygous mutation to be lost or to become fixed (Meier et al. 2014), *G* ∈ *Mat*_*N*×*K*=_({*0,1*}) is a binary matrix of genotypes per sample, and 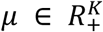 is a non-negative vector of mutation rates per genotype.

To calculate the mutational signatures across all *R* = *119* mutation types (96 single base substitutions, 2 types of multi-nucleotide substitutions, 14 types of indels and 7 types of structural variants), we used a negative binomial model to account for a higher variance compared to the mean in individual counts. For a matrix of counts *<u>Y</u>* ∈ *Mat*_*N*×*R*_(*N* ∪ {*0*}), the matrix of signatures *S* ∈ *Mat*_*R*×*K*_(*R*_+_ ∪ {*0*}) was calculated using the following model:

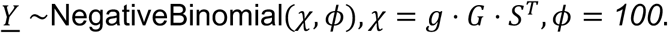

The parameter *ϕ* = *100* was based on the estimations from the data, and suggests a slight deviation from Poisson model towards higher variance. Signatures were estimated using the log-normal prior *S*_*ij*_ ∼ *logN*(*0, σ*^*2*^) with a fixed parameter *σ*^*2*^.

Mutation rates in each genotype were compared to the rates in wild-type by testing whether their squared log ratio follows chi-squared distribution. p-values were corrected for multiple testing using Benjamini-Hochberg procedure. Mutation rates per base pair per cell division were calculated assuming 15 cell divisions per generation and 2 copies of 100,272,607 bp long nuclear genome.

### Analysis of repetitive regions and G4-prone sequences

Genome-wide G4-prone sequences for *C. elegans* (Marsico et al. 2019), and repetitive regions as deposited in Repbase (www.girinst.org/downloads/repeatmaps/C.Elegans) were used to determine the association of SVs with specific genomic regions (Bao et al. 2015). For each SV, 60 bp regions around the breakpoints were overlapped with the location of G4-prone and repetitive regions. Only unique variants were used to calculate the associated proportions for each genotype. Proportions expected by chance were estimated as the ratios between the sums of all regions of interest and *C. elegans* genome size.

### Relationship to replication directionality

Directionality of replication was determined using the Okazaki fragment sequencing from (Pourkarimi et al. 2016). Fractions of Okazaki fragment reads on the minus strand 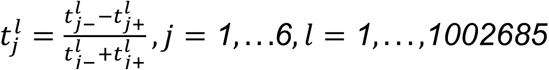 were calculated for 100-bp bins, and the bins where 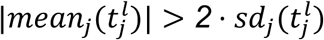 were assigned as “+” (or called right-replicating) direction if 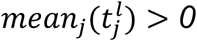, or “-” (left-replicating regions) if 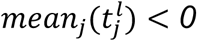. In total, we inferred the direction of replication for 45% of the genome.

### Analysis of clustered mutations

Clustering of mutations was assessed using the start points of all base substitutions and indels across samples of the same genotype and generation. Clustered status was assigned based on a hidden Markov model which predicts a series of *M* hidden states 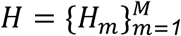, *H*_*m*_ ∈ {*clust, not*} (being in a cluster or not) for all mutations within a sample based on the distances to the next mutation 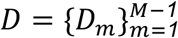, *D*_*m*_ ∈ *N* (the last mutation in each chromosome is assumed to be fixed in a non-clustered state). The probability of a set of states given the observed distances was then calculated as

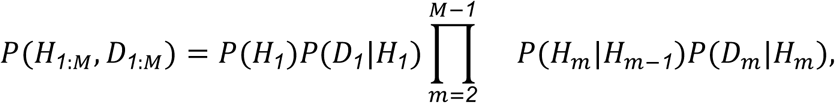

with the transition probabilities

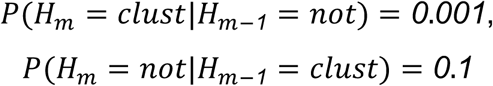

and starting probabilities *P*(*H*_*1*_ = *clust*) = *0, P*(*H*_*1*_ = *not*) = *1*.

The distribution of distances *D* given the states was assumed to be geometric, with the density of mutations within cluster assumed to be at least one mutation per 100 bases:

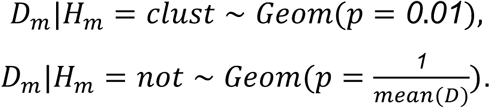

We used the Viterbi algorithm to infer the most likely set of states for each sample (Viterbi 1967).

### Microhomologies at SV and indel breakpoints

Microhomologies (MH) at the breakpoints of SVs and indels were assessed by measuring the lengths of precise alignments around each breakpoint, calculated as a sum of perfect alignment lengths between the two 30-bp regions upstream and two 30-bp regions downstream from the breakpoint sites. Only unique SVs and indels were used to calculate the proportions of variants with MH for each genotype.

## Data Access

Sequencing data and variant calling files are available under ENA Study Accession Numbers ERP000975 and ERP004086. The code for analysis of mutation rates and signatures is available on GitHub under http://github.com/gerstung-lab/mutationaccumulation.

## Acknowledgements

This work was supported by the Wellcome Trust COMSIG consortium grant RG70175, a Wellcome Trust Senior Research award (090944/Z/09/Z), a Worldwide Cancer Research grant (18-0644), and by the Korean Institute for Basic Science (IBS-R022-A1-2019) to AG. We thank the Mitani Lab funded by the National Bio-Resource Project of the MEXT, Japan, and the Caenorhabditis Genetics Center funded by the NIH Office of Research Infrastructure Programs (P40 OD010440) for providing strains. Nadezda Volkova is a member of Lucy Cavendish College, University of Cambridge. We are grateful to the COMSIG consortium for discussing unpublished data, and Kei-ichi Takeda and Orlando Schaerer for feedback and proofreading.

## Author contributions

BM, NV, AG and MG wrote the manuscript, which was approved by all authors. BM and NV analysed the data. BM, YH, SB, VGH and TP performed mutation accumulation experiments. PC and AG conceived the study. AG and MG supervised the analysis.

